# Validation of the EEG signal of the URGOnight neurofeedback device, associated with a new SMR detection method

**DOI:** 10.1101/2022.12.27.522035

**Authors:** Rudy Saulnier, Béatrice Spiluttini, Emma Touré-Cuq, Karim Benchenane

## Abstract

Sensorimotor (SMR) neurofeedback is a promising therapy for several health disorders but is still not widely used due to the high cost of the equipment. URGOnight offers a low-cost solution to democratize these therapies by providing an at-home EEG headband with dry electrodes connected to a mobile application. The first aim of this study is both to validate the URGOnight EEG signal and to compare it to Enobio-20, a medical grade EEG device. The second aim of the study is to propose a new method to detect SMR rhythm based on its oscillatory properties and discriminate it from alpha oscillations.

In our study, we compared the URGOnight headband EEG signal (C3/C4) to Enobio-20 (CP3/CP4), placed on subjects simultaneously equipped with the two headbands. All subjects (n=33) performed a dual blocking task inspired by Kulhman (1978) based on the blocking effect of movement and eyes opening on SMR and alpha respectively. This task was followed by SSVEP stimulations to evaluate the frequency response of the two EEG devices. The performance of the EEG headbands was statistically identical for most of the characteristics of the EEG signal, including the frequency response to SSVEP (from 4Hz to 20Hz). The main difference was a larger amplitude in the 8-15Hz due to the location of the reference in URGOnight that did not impair the detection of both alpha and SMR.

In addition, we show that our new method allows to discriminate alpha and SMR rhythms based on their oscillatory properties with a single recording site (C3/C4). The method is fast enough to be used in real time. We show that the detected SMR rhythm is modulated by movement as opposed to the 12-15Hz frequency band often used as indicator of SMR in most neurofeedback studies.

Altogether, our results validate the quality of the EEG recordings obtained with URGOnight since it gives similar results as the one obtained with Enobio-20, a validated EEG medical grade system. In addition, we provide a new method allowing the identification and the separation of the alpha and SMR with a single recording site C3/C4. This method opens up a new research lead to improve SMR neurofeedback efficiency and thus of its clinical possibilities by focusing on the reinforcement of the SMR oscillation strictly speaking.

**Highlights:**

- Validation of the URGOnight EEG device suitable for neurofeedback at home
- New method for the detection and the discrimination of alpha rhythm and SMR rhythm with a small number of recording sites
- The oscillatory activity related to the SMR displays different properties compared to the 12-15Hz frequency band.
- Description of a full validation procedure for wireless EEG devices usable at home for neurofeedback
- Comparison of the signal of URGOnight (dry electrodes) with a wet electrode EEG device

## 1. Introduction

### 1.1. Neurofeedback: a need for at-home EEG devices

Ongoing (spontaneous) electrical brain activity can be modified by the transient activation of specific neuronal networks in response to external stimuli (such as sensory stimulation) or internal stimuli (cognitive, such as active thinking or relaxation). Brain activity can also undergo long term changes, by modification and reorganization of neural circuits and networks in the context of learning a new skill or as a result of a pathology. This brain plasticity is at the core of neurofeedback therapies during which subjects are trained to modify their own brain activity.

Data from the literature have proven the ability of neurofeedback to improve certain cognitive aspects, such as memory or creativity (Gruzelier et al., 2014). Neurofeedback based on sensorimotor rhythm (SMR), defined by a frequency between 12 and 15 Hz, increases total sleep time (Cortoos et al., 2010; Hoedlmoser et al., 2008; Schabus et al., 2014). Neurofeedback targeting the Theta/Beta ratio, a marker of attentional drift, is used in Attention-deficit/hyperactivity disorder (ADHD) (Arns et al., 2014; Duric et al., 2012; Meisel et al., 2013; Steiner et al., 2014), and alpha waves are used in the improvement of cognitive functions (Zoefel et al., 2011), memory processes in depressive patients (Escolano et al., 2014), or to reduce migraine symptoms (Walker, 2011). These therapeutic applications are offered in a hospital setting with conventional medical systems and in the presence of a therapist. The current EEG systems utilized in the clinical practice are multichannel devices which a trained operator must position on the subject’s head and which are often cumbersome, use wet electrodes that can create skin intolerances and require hair washing after each use.

URGOnight was designed from the beginning to be a portable EEG system allowing a simple and autonomous at-home use. It operates with dry electrodes, i.e. not requiring any gel or conductive paste, and has been designed with an adjustable shape allowing a correct positioning of the electrodes by a neophyte user. This device includes the necessary channels for the collection of the SMR rhythm. An important challenge in overcoming these constraints was to maintain a quality equivalent to standard neurofeedback systems and to achieve the same EEG quality as a clinical grade system. Without a correct collection of the EEG biomarker (in this case the sensorimotor rhythm), the information sent back to the user is misleading and does not result in the desired modulation of brain signal. We will compare URGOnight to Enobio-20, a wireless medium-density EEG medical grade system for precise brain research (Ratti et al., 2017). Enobio-20 will be used as a reference system because it uses wet electrodes, which is the gold standard for high quality EEG recordings, and because it has been used to detect alpha oscillations (Vigué-Guix et al., 2020) or to study motor-related activity (Tu-Chan et al., 2017). Finally, Enobio-20 has already been used in clinical studies, for instance to detect acute pain signals from human EEG (Sun et al., 2021). **The aim of this study is to validate the quality of the EEG signal collected by URGOnight and to compare it with an electroencephalography device commonly used in clinical and research settings.**

### 1.2. Problem of identification of alpha versus SMR with two-channel EEG devices

The URGOnight EEG headband has been developed to provide at-home SMR-based neurofeedback training to help users to improve sleep quality (Krepel et al., 2021). The headband collects the EEG signal on the central C3 and C4 sites that are located above the sensorimotor cortex according to the international 10-20 classification of EEG positions on the scalp (Siuly et al., 2016) However, SMR rhythm is very close in frequency to alpha occipital waves. With classical EEG devices, the identification of these two rhythms is easily made by differentiating spatial locations of their sources: parietal for the SMR and occipital for alpha. Different techniques have been used such as spatial filtering or coherence between different recording sites (Guger et al., 2000; McFarland et al., 1997; Schoppenhorst et al., 1980). Obviously, these methods cannot be applied to devices with fewer recording sites, such as URGOnight.

For a successful sleep improvement by the SMR-based neurofeedback, it is crucial to optimize the detection of the appropriate EEG biomarker. Identifying the specific SMR frequency band for each individual and being able to distinguish it from the classical alpha oscillations could be a first promising way to improve current neurofeedback therapies. Nevertheless, taking the average of a power spectrum in a given frequency band will always return a value that does not ensure the presence of a periodical signal (referred to as true oscillation later on) in this frequency range. Thus, using a true SMR-related oscillation or an individualized value as a feedback could be a second way to improve the efficiency of SMR neurofeedback.

In order to determine individual SMR peaks and distinguish them from individual alpha frequencies, we will develop a new method to detect periodical signal corresponding to SMR and alpha (Donoghue et al., 2020). This method will be applied on EEG recordings in subjects performing a two-part protocol described in Kulhman that will be used to confirm the identity of alpha and the SMR (Kulhman, 1978). Indeed, although close in frequency, SMR and alpha rhythms are visible on the EEG when a subject is placed into different situations. The SMR is observed when the subject is immobile and decreases during movement initiation or motor imagery (Kulhman, 1978; McFarland et al., 2000). On the contrary, the amplitude of the posterior alpha waves is minimally affected by movement but is directly related to the activity in the visual cortex: alpha power increases during eye closure and decreases during visual stimulation. **Thus, the second aim of this study is to develop a new method to identify SMR and alpha rhythm as periodical signals in the EEG** (Donoghue et al., 2020) **and to use the blocking properties of SMR and alpha during the dual blocking task to validate the detection.**

## 2. Methods

### 2.1. Subjects and Data

A total of 35 individuals (18 females and 17 males; M = 25.4 years, SD = 5.3) participated in the study. One subject did not complete the entire recording session and was excluded from the study. The removed subject was a male. The second subject was excluded because of the low quality of the signal recorded (see *2.3. Data preprocessing and selection*). The remaining subjects (n = 33) included in the study had no neurological problems. The experiments were carried out in accordance with the Declaration of Helsinki. The study was approved by the Committees of Protection of Persons (Comité de Protection des Personnes ”Est IV”), declared to the French National Agency for Medicines and Health Products Safety, and the study is registered on clinicaltrials.gov. Informed consent was signed by each participant before the experiment. Subjects received monetary compensation for their participation in the experiment.

The data for the re-referencing experiment were obtained from an online database made freely available https://www.physionet.org/content/eegmmidb/1.0.0/ (Goldberger et al., 2000; Schalk et al., 2004). The dataset consists of over 1500 one- and two-minute EEG recordings, obtained from 109 volunteers. Subjects performed different motor/imagery tasks while 64-channel EEG were recorded using the BCI2000 system (http://www.bci2000.org). Only the recordings for the resting state with eyes open and eyes closed were analyzed.

### 2.2. Data acquisition with the URGOnight and Enobio-20 EEG devices

The URGOnight collects the EEG signal on the central C3 and C4 sites and reference and ground are positioned on the left and right mastoids, respectively (close to PO7 and PO8). Raw EEG signal is transmitted via Bluetooth Low Energy (BLE) technology or USB wired connection which was used in this study. Data acquisition with URGOnight was made by the software BLEexplorer. Sampling frequency was set at 500Hz for the two devices.

Enobio-20 was used as a reference system for the URGOnight. Enobio-20 device was connected to a computer via USB cable. Acquisition data with the Enobio system is made by the NIC2 software provided by Neuroelectrics. The devices were synchronized using a Transistor-Transistor Logic device (Neuroelectrics, Barcelona). Note that the systems were not designed to allow a precise synchronization that prevents a direct comparison between the signals of the two EEG devices.

### 2.3. Recording settings and behavioral tasks

The total duration of the experiment for a subject was approximately 1h30. Participants went through different tasks: dual blocking task, SSVEP, auditory oddball. For the current report, only the data recorded during the first two tasks were analyzed in this study.

Subjects were comfortably seated on a chair with armrests. The data collection was performed with both URGOnight (UrgoTech, Paris) and Enobio-20 (Neuroelectrics, Barcelona) EEG devices. The URGOnight device was installed first. The Enobio EEG device is then placed on the head with the electrodes in position CP3, CP4 and the right earlobeas reference and ground. A conductive paste (Ten20) was used for Enobio-20 electrodes. We ensured that the placement of the Enobio-20 electrodes did not disturb the position of the URGOnight headband.

#### 2.3.1. The dual blocking task

This task was inspired by Kulhman (1978) based on the blocking effect of movement and eyes opening on SMR and alpha respectively. During this task, participants followed the protocol: 1) 1 minute, eyes open, immobile; 2) 1 minute, eyes open while performing bilateral movements of forearms, wrists and hands; 3) 1 minute, eyes closed, immobile; 4) 1 minute, eyes closed while performing bilateral movements of the forearms, wrists and hands.

#### 2.3.2. SSVEP task

The intermittent light stimulation device (PandoraStar stroboscopic lamp) was placed at a distance of 1.10m from the subject on a stand at 3 feet height. The intensity level of the lamp was set at 5. The stroboscopic lamp was switched on with a rhythmic stimulation mode. Each stimulation was 1 minute long at the frequencies of 4Hz, 8Hz, 10Hz, 13Hz, 15Hz or 20 Hz. During the whole protocol, the subjects seated comfortably while keeping their eyes closed to limit artifacts in the signal. The task starts with 1 minutes of resting state (immobile/eyes closed). Then they were stimulated 6 times (corresponding to each frequency used) for 1 minute with 1-minute inter-stimulation interval.

### 2.4. Data preprocessing and selection

The preprocessing steps were identical for both systems. Obtained data were bandpass filtered between 1Hz and 40Hz with numerical Butterworth filter of order 4. Signal was considered as an artifact when its amplitude exceeded 50µV, if its kurtosis was outside of the chosen boundaries of 1-6, or if the power of the low-pass filtered signal at 6Hz surpassed 20 µV². Artifacts were removed by epoch of 1s. The entire session was removed from the dataset if this procedure resulted in keeping less than 20% of the original signal.

### 2.5. Data analysis and spectral analysis

Data analysis was performed with custom Matlab scripts (Mathworks, Natick, MA, USA). The signal was translated in time-frequency domain by iteratively calculating the power spectra on a time bin of 3 sec with 100 ms long step using the mtspecgramc.m function (tapers [2,3]) from the chronux toolbox (http://www.chronux.org). The spectra were calculated for a frequency range of 0-40 Hz, with a sampling frequency of 500Hz. Signal to noise ratio (SNR) was performed as in the study by Kam and colleagues. (2019).

### 2.6. Detection of bump in the power spectrum and histogram

Periodic signal will lead to a bump in the power spectrum, on the contrary to aperiodic signal associated with a 1/f decay. To detect bumps in power spectra (or histograms), we used the method developed by Donoghue and colleagues (2020). Briefly, this method aims at modelling the signal into non-periodic and aperiodic components, with no a priori specification of frequency band. Practically, this method relies on the identification of several bumps in a power spectrum after removing the exponential decay of the spectrum (1/f component). In order to increase the speed of the algorithms, we replace the fitting procedure used in the original publication by the removal of the 1/f component of the power spectrum obtained by multiplying each power spectrum by the frequency.

We identified bumps in the power spectra (after removal of 1/f component) or histogram, by using the following procedure: 1) Define a threshold for the detection of peak in the spectrogram: mean + 2 standard deviation, 2) Detect the max of the power spectrum corresponding to the peak of a bump. 3) Define the two values corresponding to 2/3 of the max values as the width of the bump and select the one closest to the peak. 4) Define a gaussian with the peak of the bump as a mean, with a sigma computed to match the width obtained in step 3. 5) Remove the gaussian from the original signal. 6) Apply this method again to detect other peaks.

In order to identify both alpha and SMR, we used the method described above and calculated the histogram of all oscillation frequencies detected in the 2-20Hz frequency band over the entire double blocking task. We then force the identification of two bumps in the histogram with the method used for the power spectra but without any threshold. These two bumps were fitted with two gaussians of which the intersection determined the frontier between SMR and alpha. According to the literature, alpha is most frequently observed, with higher power, compared to SMR (Kulhman, 1978). Therefore, the largest hump detected in the histogram, which corresponded to the most frequently detected oscillations, was defined as alpha, with the second hump (if detected) corresponding to the SMR. Each time bin of the spectrogram is then re-analyzed to detect bumps, with the 2 SD threshold, and in the 4 possible classes: no oscillation, SMR, alpha or SMR+alpha. The power spectra of all periods belonging to each of the 4 classes were averaged (note that the previous step with the histograms ensured to observe a peak in this averaged power spectrum). The frequency of SMR (or alpha) is given by the peak of these averaged power spectra. Then the time bins for the alpha and SMR were redefined according to this frequency range. The amplitude of the bumps for all these time bins was averaged to determine the amplitude of the rhythm. The described steps resulted in a representation of each type of the oscillations as a percentage of time during which the particular rhythm has been detected and the amplitude of each rhythm detected in the spectrogram.

### 2.7. Statistics

Statistics were performed with JASP, an open-source program for statistical analysis supported by the University of Amsterdam (https://jasp-stats.org). For most of the analyses, we used ANOVA with repeated measures and several types of post-hoc tests were specified when used. Effect sizes were computed with the Cohens d, with values d = 0.2, 0.5 and 0.8 corresponding respectively to small, medium and large effects.

For comparison of the EEG devices, we used Bayesian statistics. Bayesian framework provides an experimenter with means to quantify the degree of similarity, in opposition to the conventional statistical test that tests the difference. The equivalent of the p-value for this type of test is the factor Log(BF_10_). If this factor is negative, the two groups cannot be considered identical. If it is positive, the two groups are considered identical. The level of evidence are: from 0 to 0.48: ns, from 0.48 to 1 : anecdotal evidence, from 1 to 1.48 : moderate evidence, from 1 to 1.48 : strong evidence, from 1.48 to 2 : very strong evidence, >2 : extreme evidence (Jeffreys, 1961).

## 3. Results

A total of 35 participants participated in the study. One subject did not complete the entire recording session and was excluded from the study. The second subject was excluded because of the low quality of the signal recorded. Thus, analysis was performed on 33 remaining participants (17 females and 16 males; M = 25.2; SD = 5.2). At the stage of the automatic data selection (see methods), 86% of the signals was selected for the Enobio-20 and 80% for the URGOnight EEG device. The same procedure can be applied to the data acquired by the URGOnight (Fig 1A) during at-home neurofeedback sessions to ensure quality of the signal recorded by a non-expert user.

**Figure 1.**
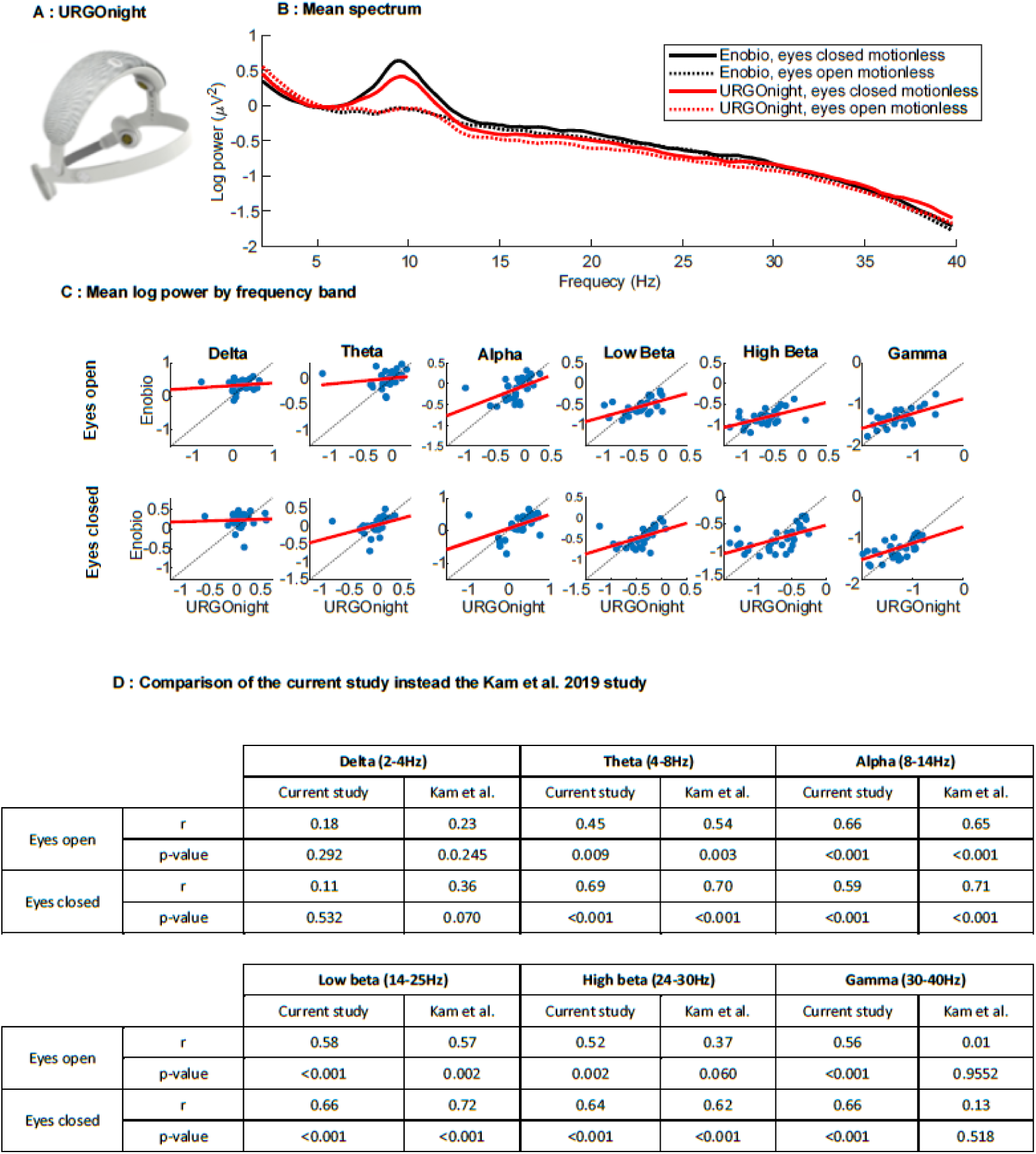
Comparison of the broadband signal between URGOnight and Enobio. **A.** URGOnight headband, adjustable to head size, with two measuring dry electrodes over the sensorimotor cortex C3 and C4 with references close to PO7. **B.** Power spectrum of the signal of the two EEG devices for rest with eyes open and eyes closed in log scale. Black: URGOnight. Red: Enobio. C. Correlation of the spectral power for selected frequency bands recorded with the two headbands. Delta: 2-4Hz; Theta: 4-8Hz; Alpha: 8-14Hz (as in Kam et al., 2019); Low beta: 14-24Hz; High beta: 24-30Hz; Gamma: 30-45Hz. **D.** Comparison of the correlations between the power in each frequency band in our study and in a reference study that aimed at comparing two EEG devices with dry and wet electrodes (Kam et al., 2019).

### 3.1. Comparison of the broadband signal between URGOnight and Enobio

The protocol for this study was largely inspired by the one used in the article by Kam and colleagues (2019), which compares EEG devices with dry and wet electrodes. The wet electrodes are used as a gold standard. Since URGOnight has dry electrodes and Enobio has wet electrodes, we performed the same analyses as in the study by Kam and colleagues (2019) in order to compare URGOnight (dry electrodes) to the signals obtained with Enobio (wet electrodes), a validated wireless medium-density EEG medical grade system.

We first analyzed the EEG obtained with the two EEG devices in immobile subjects with their eyes open or closed. The power spectrum density of the two systems was very close across the entire frequency range shown (Fig. 1B). We then performed correlation of the mean power for each system in different frequency bands and compared the results with the ones obtained in the study by Kam and colleagues (2019) (Fig 1C, Table 1). For both eyes open and eyes closed conditions, the mean absolute power of theta, alpha, and low beta positively correlated between URGOnight and Enobio across subjects (r > 0.54, p < .005). Generally, the correlation coefficients were very close to the ones observed in the study by Kam and colleagues. As in Kam et al (2019), the only non-significant correlation was found for delta frequency. The power differences observed in the lower frequencies cannot be attributed to differences in signal-to-noise ratio (SNR) (SNR in the delta range: z = 1.54, p = 0.12, SNR in the theta range: z = 0.42, p = 0.67, Global SNR: z = 0.37, p = 0.71). The origin of the difference between the two devices in the low frequency is still unclear. Thus, except for the delta frequency band (2-4Hz), our results confirm that the data acquired with dry (URGOnight) and wet (Enobio-20) electrodes are close to identical in the same conditions.

**Table 1.** Detection of Alpha and SMR

| N=33 | Alpha | SMR | Alpha & SMR | Alpha/noSMR | noAlpha/noSMR | SMR/noAlpha |
| --- | --- | --- | --- | --- | --- | --- |
| *Urgo* | 32 (97%) | 22 (66%) | 22 (66%) | 10 (30%) | 1 (3%) | 0 (0%) |
| *Enobio* | 30 (91%) | 28 (84%) | 28 (84%) | 2 (6%) | 3 (10%) | 0 (0%) |

### 3.2. Comparison of the raw alpha band signal between URGOnight and Enobio

We then compared the maximum power in the alpha band (8-12Hz) between the two EEG devices. Subjects performed 3 times the eyes open/immobile task throughout the whole experiment. Power in the alpha band was thus quantified during these three 1-minute-long sessions in order to test the stability and consistency of the recordings obtained with URGOnight and to compare them with those obtained with the Enobio-20 reference headband.

In the averaged power spectrum, we clearly see a peak around 10Hz (Figure 2A). The values of the peak for both URGOnight and Enobio give similar results for the three different sessions (Fig 2B, two-way ANOVA with repeated measure, main effect repetition, F(2,64)=0.70, p=0.50), Log(BF_10_)= −2.738****). The frequency distribution of the peaks in the alpha band were similar and also reproducible (Fig 2C), two-way ANOVA with repeated measure, main effect EEG device: F(1,32)=1.353, p=0.253, Log(BF_10_)= −0.387, main effect repetition: F(2,64)=1.449, p = 0.242, Log(BF_10_)=-2.101****. Subsequent Bayesian post-hoc test confirms that the frequencies come from the same distribution (Log(BF_10,U_)=−0.946*). Altogether, this shows that within an EEG device, the characteristics of the largest oscillation in the 8-12Hz frequency band are reproducible across different recording sessions.

**Figure 2.**
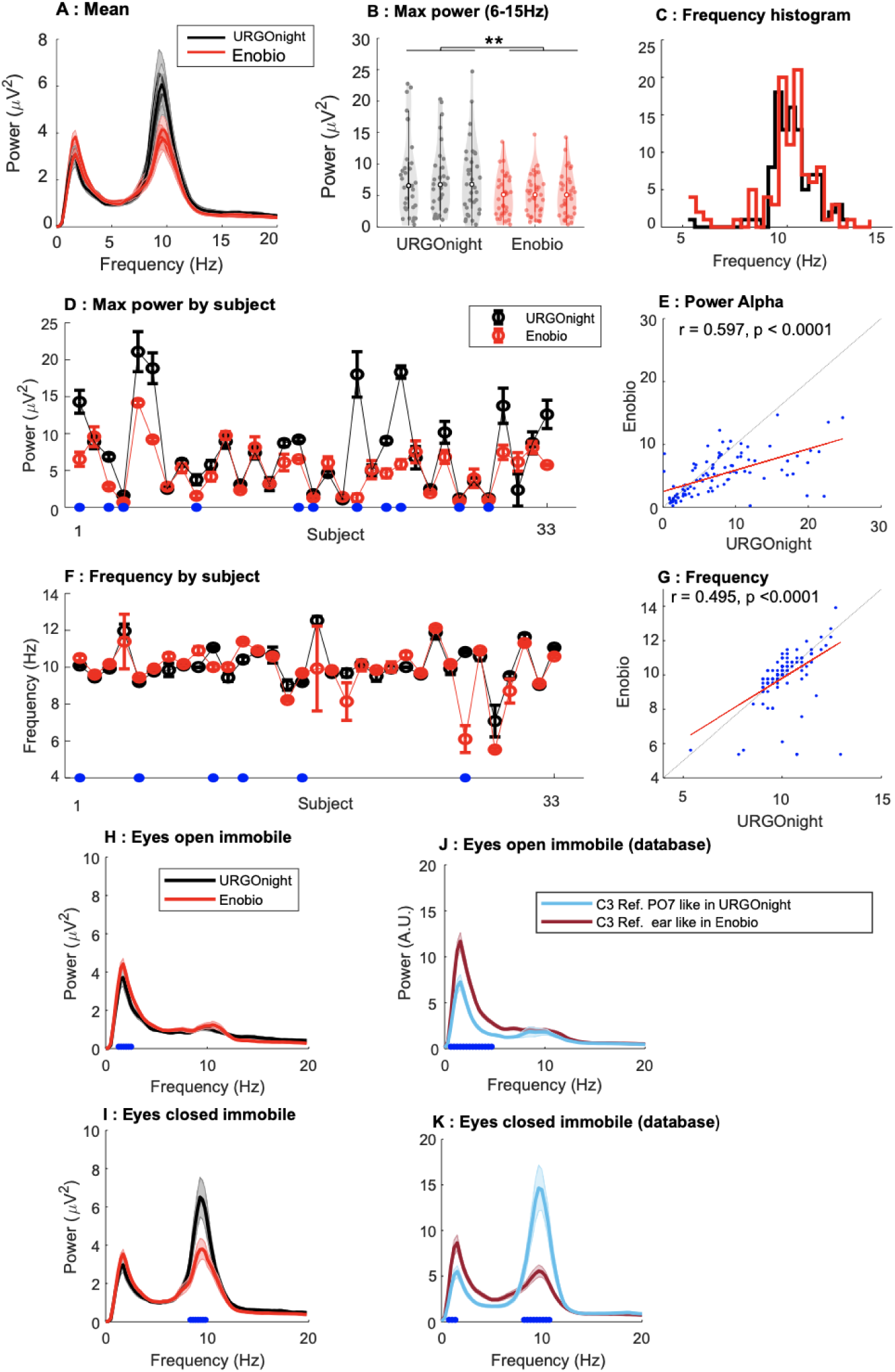
Comparison of EEG headbands. Black: URGOnight URGOnight, Red: Enobio. **A.** Average of the 3 eyes closed recording session for each EEG device. mean+/−SEM. Quantification of the maximal power in the alpha band during the 3 eyes-closed periods for the 2 EEG devices. Two-way ANOVA with repeated measures: main effect EEG device, F(1,32)=0.578, p=0.002, main effect recording session show a statistical equality, F(2,64)=0.70, p=0.50, Log(BF_10_)= −2.738****. **C.** Distribution of the frequency for the maximal power in the alpha band. Two-way ANOVA with repeated measures: F(2,64)=1.449, p=0.242, Log(BF_10_)= −0.387, Log(BF_10,U_)=−0.946*, and different sessions (Fig 1B) are statistically identical too (F(2,64)=1.449, p = 0.242, Log(BF_10_)=-2.101****). **D,F** Average power and frequencies for the maximal power in the alpha band detected for individual subjects. *Blue dot*: significant difference between the EEG devices, Post-Hoc Fisher test p<0.05. **E,G** Correlation between amplitude and frequency of the maximum peak recorded in eyes close condition. Correlations are both significant (power: r=0.60, p<0.0001, n=99, frequency: r=0.49, p<0.0001, n=99). **H-K.** Effect of the position of the reference electrode on the maximum peak in the alpha band. HI, Power spectrum in eyes close and eyes open condition in our study (**H,I**, black URGOnight, red Enobio). **J,K**. Same as H-I but with data from a database made available by Schalk et al. 2004 with a modification of the reference to match the URGOnight EEG device. This database contains 109 subjects performing the eyes - open motionless task for 1 min and the eyes-closed motionless task for 1 min. *Dark red*: average of the 109 subjects with the ear reference as for the Enobio headband in our case. *Light blue*: average of the 109 subjects after subtracting from the C3 electrode the PO7 electrode to match the recordings obtained with URGOnight. Note the large increase in power in the alpha band induced by the re-referencing procedure that reproduces the difference observed between URGOnight and Enobio.

This analysis also shows that the amplitude of the peak in the alpha band was larger for URGOnight than for Enobio (Fig 2B, two-way ANOVA with repeated measure, main effect EEG device, (F(1,32)=0.578, p=0.002). The analysis across individual subjects reveals that the increase in the alpha power with URGOnight is restricted to a subpopulation of subjects. Only 11 subjects out of 33 show a significant difference of the maximal power in the alpha band. The difference between the two EEG devices is important in only four of these subjects (Fig. 2D-G). Moreover, the difference in frequency is extremely small and reaches significance only for 6 subjects out of 33. The individual values of both amplitude and frequency of the maximal peak in the power spectrum are significantly correlated (Fig. E,G - power: r=0.60, p<0.0001, n=99, frequency: r=0.49, p<0.0001, n=99).

### 3.3. Origin of the increase power in unprocessed alpha band with the URGOnight device

We then investigated the origin of the increased alpha power in URGOnight recordings. As described previously, URGOnight was designed from the beginning to be a portable autonomous EEG system for the non-expert users. It has an adjustable shape that promotes correct positioning of the electrodes even by a neophyte user. The tradeoff between usability and the quality of the signal lead to the compromises in the actual design where recording electrodes were placed in C3 and C4 positions, and reference and ground close to PO7 and PO8. Enobio-20 recording electrodes are located in CP3 and CP4 with a reference attached to the right ear.

In preliminary experiment, we observed no difference between the signal obtained in C3 position (URGOnight) and CP3 (Enobio-20), consistent with the literature. Yet the position of the reference could affect the amplitude of the peak in the alpha band. We investigated this possibility by using an open database of EEG recordings done in 109 subjects placed in eyes open-eyes closed conditions (Goldberger et al., 2000; Schalk et al., 2004). Using this database, we compared the recordings obtained in C3 with the reference on the ear (as for Enobio-20) and with the reference located at PO7 (as for URGOnight). Strikingly, the recordings with the reference at PO7 position demonstrated a large increase in the amplitude of the peak in the alpha band, similar to what was observed with URGOnight (Fig 2K). In addition, similar to the URGOnight recordings, alpha peak vanished in the eyes open condition suggesting that is comes from the occipital alpha (Fig 2H,J). These results point at the fact that the difference in alpha power observed between URGOnight and Enobio recordings is caused by differences in the location of the reference electrode and comes predominantly from the occipital alpha.

### 3.4. Validation of the frequency response of the devices with SSVEP

We then decided to further confirm that the difference in the amplitude of the peak in the alpha band observed in URGOnight is not due to a problem in the frequency-specific response of the device. In order to rule out this possibility, we used SSVEP protocol to compare the frequency response of the EEG devices with a minimum of variation due to endogenous oscillations (that could be specific to each subject). Indeed, exposition to the rhythmic visual stimulus generates controlled oscillations at the chosen frequency in the visual cortex. SSVEP protocol guarantees the presence of oscillations at the specific frequency generated in the particular brain structure, which altogether creates a highly controlled setting to study oscillatory phenomena.

The SSVEP task is composed of a 1-minute long period with eyes closed, followed by 6 periods lasting 1 minute with SSVEP stimulations of 4, 8, 10, 13, 15 and 20Hz as described in the methods section. Oscillations generated by SSVEP are not stable and can vary with attention (Morgan et al., 1996). Thus, we selected periods with SSVEPs in the EEG when the coherence between the light stimulation and the EEG signal is higher than 0.9.

We can see clear peaks corresponding to the SSVEP stimulation for the two EEG devices regardless of the SSVEP frequency. The power spectrum reveals a peak at around 10Hz, that is reminiscent of the average spectrum visible during the closed eyes condition. We measured the amplitude of the power spectra and observed that the amplitudes of the responses to the SSVEPS were larger for the URGONight headset at frequencies of 10Hz and 13 Hz, an effect that is coherent with the previous observations. In order to control for effect observed in resting state, we quantified the response to the SSVEPs by removing the baseline. We computed the baseline by taking the power spectrum during resting state with eyes closed without any stimulation. After this correction, the response of both EEG devices became almost flat and the resonance observed around 10Hz vanished. Two-way ANOVA with repeated measures yields no main effect of the EEG device (F(1,32) = 0.919, p = 0.345, Log(BF_10_) = −1.713***). There is a main effect of the frequency of the SSVEP stimulation (F(5,160) = 2.270, p = 0.05, Log(BF_10_) = 0.989*), but that is mainly due to the 20Hz frequency and not to the resonance at 10Hz as observed previously (F(4,128) =1.162, p=0.331 after removing the 20Hz condition). This shows that the response of URGOnight is similar to the one observed with Enobio-20 for every frequency ranging from 4 to 20Hz. The two-way repeated measures ANOVA also reveals that SSVEP was slightly more frequently detected with URGOnight compared to Enobio (main effect of the EEG device: F(1,32) = 4.985, p = 0.033).

Together with the previous results, this shows the electrical signal obtained with URGOnight is similar to the one observed with Enobio with an identical frequency response (between 4Hz and 20Hz), and that the difference observed in figure 3C is most likely due to the position of the reference.

**Figure 3.**
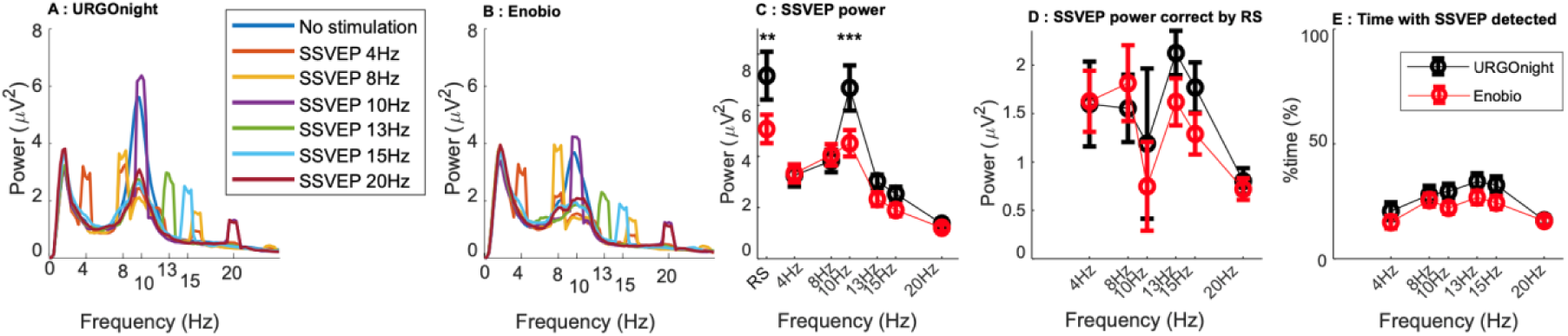
Comparison of SSVEP between URGOnight and Enobio. **A,B**. Average spectrum of the 33 subjects for the SSVEP protocol for the URGOnight EEG headband in **A** and Enobio in **B** with the C3 and CP3 electrodes respectively. **C.** Comparison of the amplitudes of the response to SSVEP stimulations for the two EEG devices (URGOnight: *black*, Enobio: *red*). The amplitude is defined as the maximum power at the SSVEP stimulation frequency ±0.6Hz. The statistical analysis used was a repeated measure two-way ANOVA: main factor EEG headbands (F(1,32) = 12.437, p = 0.001) and main factor SSVEP frequency (F(6,192) = 20.521, p<0.001) and a strong interaction F(6,192) = 4.879, p = <0.001). Post-HOC Holm test, showed significant differences between the two headbands for the 10Hz SSVEP stimulation (p < 0.001) and for resting state situation (p = 0.001**). **D.** Comparison of the amplitudes of the SSVEP stimulations after correction from the baseline (two-way ANOVA with repeated measures, main factor EEG device : F(1,32) = 0.919, p = 0.345, Log(BF_10_) = −1.713***, main factor SSVEP frequency : F(5,160) = 2.27, p = 0.050, this effect only depends on the 20Hz stimulation see Text). Note the absence of resonance at 10Hz after renormalization. **E.** Percentage of time during which SSVEP is detected as a function of EEG headband and frequency. The two way repeated measures ANOVA gives a difference between the two headbands (F(1,32) = 4.985, p = 0.033). There is a main effect for frequency (F(5,160)=6.582, p<0.001) but no interaction F(5,160) = 0.988, p = 0.427). All percentages have been corrected in arcsin of the square root of the percentage before ANOVA.

### 3.5. Behavioral task for alpha and SMR segregation

An accurate simultaneous detection of both the SMR and alpha is made difficult because alpha oscillations prevent direct observation of the SMR due to significant overlap in their frequency bands. The situation is further complicated by the fact that the frequency bands of the SMR and alpha could significantly and independently vary across subjects (Kulhman, 1978). We therefore used the dual blocking task inspired by Kuhlman (1978) to dissociate alpha and SMR rhythms (and allow them to be separated). When the subject is immobile, the SMR (parietal) rhythm increases and the oscillation is suppressed by the motor activity. Similarly, the occipital alpha oscillations emerge when the subjects close their eyes and disappears in the eyes open condition. With this procedure, we can identify the four patterns of oscillatory activity (Figure 4): 1) presence of alpha and the SMR in immobile subject in eyes closed condition, 2) presence of alpha alone in a moving subject with eyes closed, 3) presence of the SMR alone for an immobile subject in eyes open condition, and finally 4) absence of any oscillation in a moving subject with open eyes.

**Figure 4.**
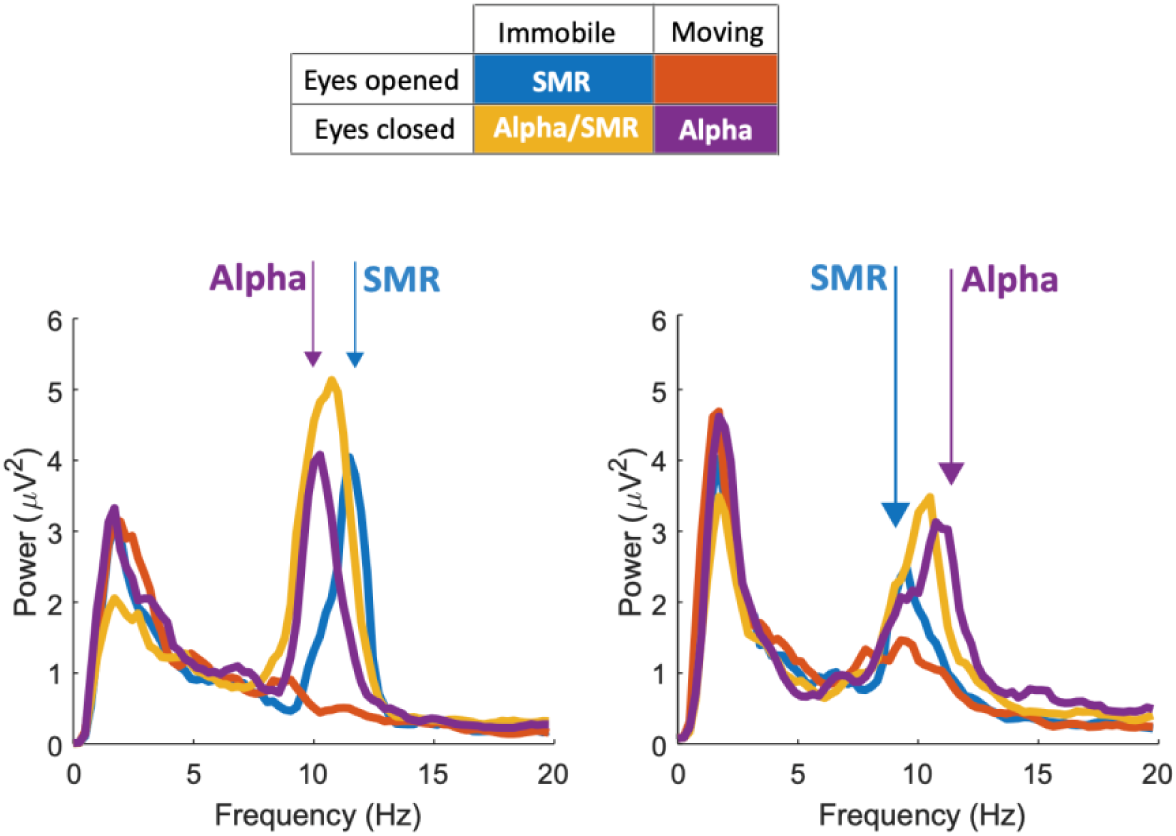
Example of spectrogram obtained in the dual blocking task used to dissociate the SMR and alpha. *Top,* Description of the experimental task with conditions: 1) presence of alpha and the SMR in immobile subject with eyes closed (yellow), 2) presence of alpha only in moving subject with eyes closed (purple), 3) presence of SMR only for immobile subject with eyes open (blue), and finally 4) absence of any oscillation in moving subjects with open eyes (red). Bottom, average power spectrum for two subjects in the experimental situations with URGOnight. The two subjects both show two separate peaks in the spectrum that exhibit the functional characteristics of the occipital alpha rhythm and of the SMR.

In Figure 4, we can see two examples we obtained with the URGONight device. This procedure allows an independent manipulation of the two types of oscillations that are identifiable in the average power spectrum for each situation, reproducing the results obtained in the study by Kuhlman (1978). However, this effect is not clearly identifiable in all individuals. Moreover, the identification of the two peaks based on the average spectrogram is difficult to automatize in most cases. In order to use URGONight for at-home SMR-based neurofeedback sessions, we therefore need a robust and reliable automatic method to separate the SMR from the alpha in real time.

### 3.6. Method of segregation between alpha and SMR

The different situations of the dual blocking task will induce the independent emergence of alpha and the SMR. Our approach was then to identify two types of oscillations in the spectrogram without any a functional a priori and then to verify whether the two oscillations exhibit the functional characteristics of the occipital alpha rhythm and of the SMR.

To create an automated technique for SMR-alpha differentiation, we set-up to identify the two types of oscillations without any a priori knowledge about the conditions, during which the data were recorded, and then to verify that the signal in the obtained frequency bands satisfy the definition of the SMR or alpha oscillations. In order to detect different types of co-occurring oscillations, we used the method developed by Donoghue and colleagues (2020). Briefly, this method aims at modelling the signal into non-periodic aperiodic components, with no a priori specification of the frequency band. Practically, this method relies on the identification of several bumps in a power spectrum after removing the exponential decay of the spectrum (1/f component).

Although the method developed by Donoghue and colleagues (2021) worked well in our data, the program is computationally heavy since it relies on fitting an exponential decay distribution at each time bin of the spectrogram. In order to increase the speed of the algorithms, we replaced the fitting procedure used in the original publication by th e removal of the 1/f component of the power spectrum obtained by multiplying each power spectrum by the frequency. We found the same results with the two methods but the latter was considerably faster and thus could be used in real-time neurofeedback application.

For each temporal bin of the spectrogram, peaks in the power spectra with removed 1/f component were detected using a threshold of mean power plus two standard deviations. Afterwards, we have built the histograms of the instantaneous frequency for the two most powerful oscillations over the entire dual blocking task (Fig 5H). The histogram indicates the percentage of time for which an oscillation with a given frequency is detected. According to the literature, alpha is a more prominent and more frequently observed type of oscillations compared with the SMR (Kuhlman 1978). For that reason, we identified the largest bump detected in the histogram (i.e. the most frequently detected oscillations) as alpha, and the second most frequent bump (if detected) corresponded to the SMR. Globally, this method tells us for each time bin whether we are in one of the four possible situations: 1) absence of any oscillation 2) presence of alpha, or 3) presence of both alpha and 4) SMR with their respective amplitude.

**Figure 5.**
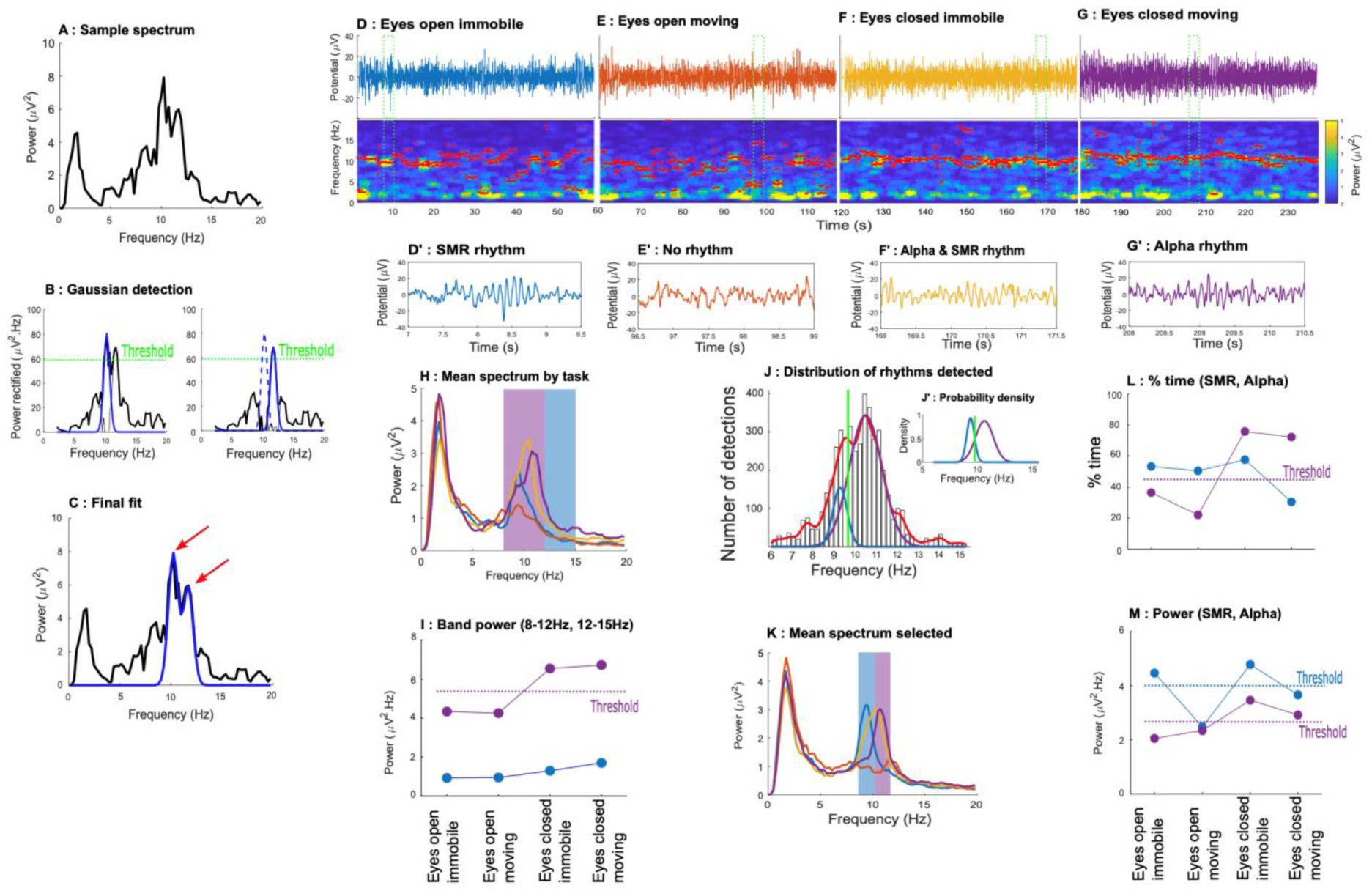
Method of detecting Alpha and the SMR brain rhythms. **A**. Example of a spectrum between 0Hz and 20Hz with the URGOnight headband (electrode C3). This spectrum has three peaks: one at about 2Hz and two others between 10 and 12Hz. The peak at 2Hz corresponds to the ~1/f decay altered by the high pass filter as shown by Donoghue and colleagues (2021). The other two peaks likely correspond to the Alpha rhythm and/or the SMR rhythm. **B.** Correction of the power spectrum to remove the 1/f component. In order to correct for the ~1/f decay, the spectrum is multiplied by the frequency, cut between 2Hz and 20Hz and the minimum value is removed. A threshold is set to the average of the spectrum + 2 standard deviations. Then by iteration, Gaussians are removed from the spectrum until the amplitude of the spectrum is below this threshold. This technique is similar to that described in Donoghue et al. **C.** Blue line corresponds to the sum of the two detected gaussians. **D-G**. Example of the application of the method for the dual blocking task consisting of 4 parts. 1st line: raw signals. 2nd line: spectrogram. The red points correspond to the average of the detected gaussians. 3rd line: 2.5s of raw signals selected in the green frames. **H and I.** Unprocessed average power spectrum and evolution of the power in the 8-12Hz (purple) and 13-15Hz (blue) in the dual blocking task : 1) presence of alpha and SMR in immobile subject with eyes closed (yellow), 2) presence of alpha only in moving subject with eyes closed (purple), 3) presence of SMR only for immobile subject with eyes open (blue), and finally 4) absence of any oscillation in moving subjects with open eyes (red). Note that the movement does not modify the power in the 13-15Hz frequency range. **J.** Distribution of the average frequency of the detected gaussians between 6Hz and 16Hz. The histogram of this distribution is estimated by the Kernel method. This density is split into two Gaussians. The higher one is considered as the Alpha rhythm and the second as the SMR rhythm. (insert: J’ Separation of the two Gaussians). **K.** Average of the spectrogram for the periods for which alpha (purple) and SMR (blue) are detected, when the two rhythms are detected (yellow) and when no oscillation is detected (red). **L.** Percentage of time during which the rhythms are detected (alpha: purple and SMR: blue). For alpha, for instance, closing the eyes increases the time with an alpha rhythm as expected, but the percentage is not modified by movement. **M.** Same as in L but for the amplitude of the alpha and SMR when the oscillation is detected. Note the decrease of alpha after eye opening and the decrease of SMR with movement.

Importantly, in this example, the oscillations that we defined as alpha exhibits the functional characteristics of the classical occipital alpha rhythm: it showed a decrease in the power during eyes open condition (Figure 5H in purple). Similarly, the oscillation defined as the SMR also exhibited a decrease in power during active movement (Figure 5H in blue). It is worth pointing out that although the quantification of alpha as the averaged power in the 8 - 12Hz frequency band behave as expected for visual alpha rhythm, the SMR defined as the averaged power in the 12-15Hz (a method often used in SMR neurofeedback studies) failed to demonstrate functional characteristics of the SMR (see fig. 7 for a quantification on the entire dataset).

### 3.7. Comparison of alpha and SMR between URGOnight and Enobio

Using the method described above, we were able to detect alpha in 97% (32/33) of the subject with URGOnight and in 91% (30/33) of the subject with Enobio (Table 2). The SMR was found in 66% (22/33) of the subject with URGOnight (alpha vs SMR, Khi2=10.1852, p=0.0014) and in 84% (28/33) of the subject with Enobio (alpha vs SMR, Khi2=1.4383, p=2304). Thus, the detection of both alpha and the SMR was similar with the two EEG devices, even if the SMR was slightly more difficult to detect with URGOnight (but this difference did not reach statistical significance level, SMR: Khi2=2.9700, p=0.0848, alpha: Khi2= 0.3492, p=0.5546). Note that the detection of alpha and SMR in our study are very close to what observed in Kuhlman (1978).

**Table 2.** 3-way ANOVA: 1) EEG device, 2) Eyes 0/C, 3) Movement/Immobility.

|  | Alpha Power, n=26, df = 1 |  | Alpha (%time), n=26, df = 1 |  | SMR Power, n=19, df = 1 |  | SMR (%time), n=19, df = 1 |  |
| --- | --- | --- | --- | --- | --- | --- | --- | --- |
| EEG device | $F=2.25$ | $p=0.14$ | $F=0.35$ | $p=0.56$ | $F=10.53$ | $p=0.004^{**}$ | $F=1.33$ | $p=0.26$ |
| Eyes | $F=126.45$ | $p<0.001^{****}$ | $F=120.73$ | $p<0.001^{****}$ | $F=32.66$ | $p<0.001^{***}$ | $F=2.29$ | $P=0.14$ |
| Movement | $F=13.26$ | $p=0.001^{***}$ | $F=2.89$ | $p=0.1$ | $F=32.75$ | $p<0.001^{****}$ | $F=7.89$ | $p=0.01^{**}$ |

|  | Power 8-12Hz, n=33, df = 1 |  | Power 12-15Hz, n=33, df = 1 |  |
| --- | --- | --- | --- | --- |
| EEG device | $F=11.28$ | $p=0.002$ | $F = 15.246$ | $P<0.001^{***}$ |
| Eyes | $F=72.34$ | $p<0.001$ | $F = 16.361$ | $p<0.001^{***}$ |
| Movement | $F=18.42$ | $p<0.001$ | $F=7.526$ | $p=0.01^{**}$ |

The frequencies of the SMR and alpha detected by the two EEG devices are highly correlated (r=0.852, r=0.852, p<0.001; r =0.834, p<0.001). We used Bayesian statistics to show that the frequencies of the two rhythms are statistically identical for both EEG devices (SMR: Bayesian significance=5611, extreme evidence, alpha: Bayesian significance=1.3e6, extreme evidence). The distributions for both SMR and alpha rhythms are identical with a mean around 10Hz, but the SMR rhythm has a greater dispersion (SMR-URGOnight = 10.81+/−0.47, SMR-Enobio = 9.68+/−0.35; alpha-URGOnight = 10.08+/−0.16, alpha-Enobio = 10.17+/−0.15; Wilcoxon test for SMR: W=0.881, n=19, p=0.1, Log(BF_10_) = 0.198 (ns), and for alpha: W=0.946, n = 30, p=0.1, Log(BF_10_) = −0.066 (ns)).

**Figure 6.**
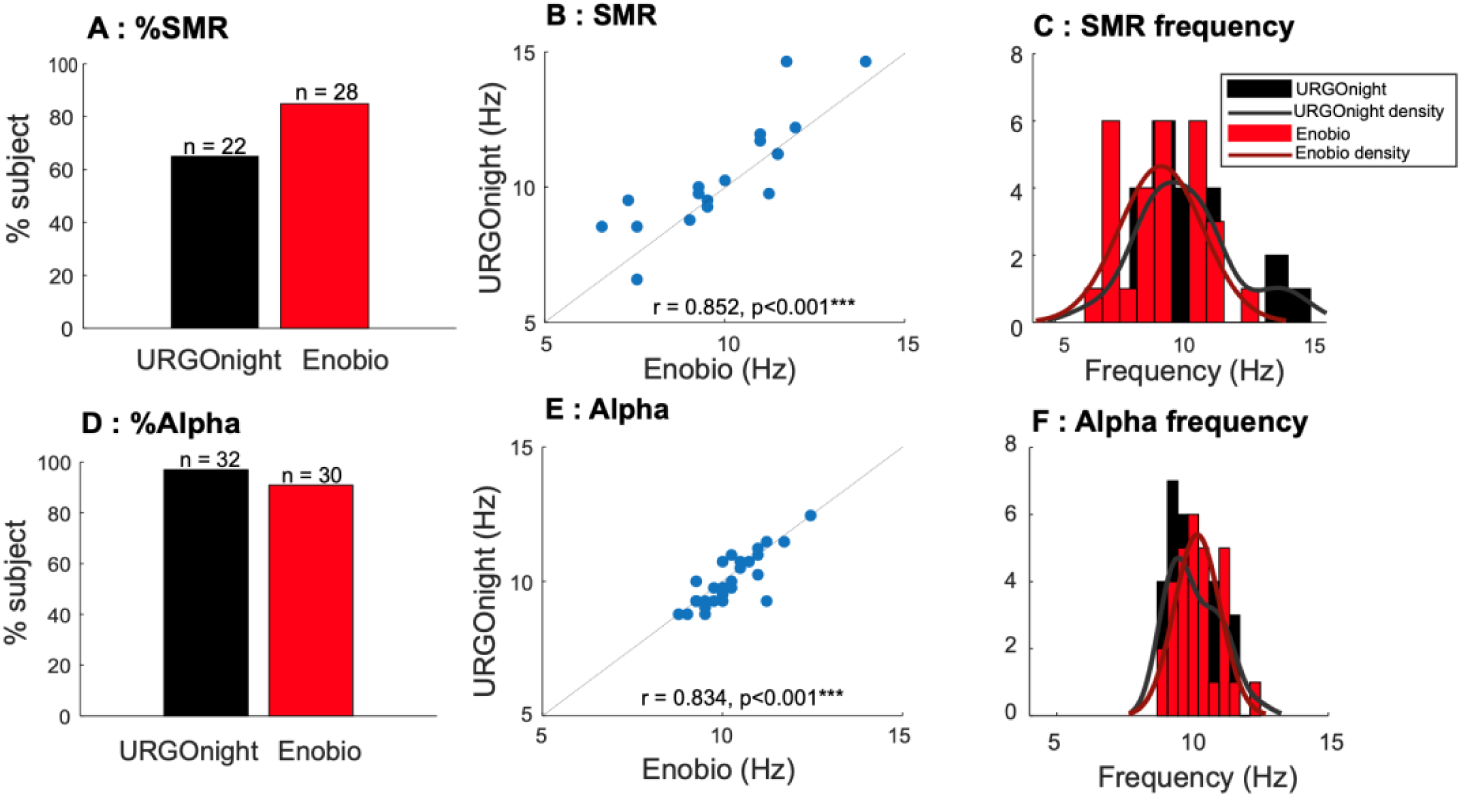
Distribution of Alpha and SMR detected by our method. **A,D**. Percentage of subjects for which each rhythm was detected. **B,E**. Pearson correlation between the frequency of the detected rhythms obtained with the two EEG devices. **C,F**. Distribution of the frequencies of the rhythms detected by our methods with the two EEG devices. The distributions for both SMR and alpha rhythms are identical with a mean around 10Hz (Wilcoxon test, SMR W=0.881, n=19, p=0.1, Log(BF10) = 0.198 (ns), alpha W=0.946, n=30, p=0.1, Log(BF10) = −0.066 (ns)).

The method of rhythm identification used in the current study relies only on the frequency domain of the oscillations and does not use any a priori knowledge about the functional properties of the detected alpha and SMR. We analyzed the effect of the different experimental conditions of the dual blocking task on the SMR and alpha. As in Kuhlman (1978), we expect a decrease in the detected SMR rhythm when subjects move and a decrease of the alpha rhythm when the subjects open their eyes.

As expected for alpha, the power of the oscillation detected as alpha decreased in almost all the subjects when in the eyes closed condition for the two EEG devices: 94% (30/32) with URGOnight and 100% (30/30) with Enobio. This was done in moving subjects, but the same result is observed for immobile subjects.

The power of the oscillation detected as the SMR (in the eyes open condition) decreases during movement in 76% (16/21) with URGOnight and in 78% (22/28) with Enobio (URGOnight vs Enobio: Khi2= 0.0391, p=0.8433). In the eyes closed condition, the oscillations detected as the SMR exhibited a decrease in power during movement in 81% (17/21) of the subjects with URGOnight and in 86% (24/28) of the subjects with Enobio (URGOnight vs Enobio: Khi2= 0.1992, p=0.6554).

Notably, if we define the SMR as the average power in the 12-15Hz frequency band, the effects are almost at the chance level: 48% (16/33) for URGOnight and 54% (18/33) for Enobio (Our method vs. 12-15Hz: Khi2= 4.0803, p=0.0434for URGOnight, Khi2= 3.8733, p=0.6554 for Enobio). Results are confirmed if we quantify the 12-15Hz only in subjects for which of the SMR detection method was effective.

As described previously, our method allows us to identify the periods during which we detect a given rhythm but also the power of the oscillations when the rhythms are detected. The quantification for the two parameters is shown in figure 7. For percentage of time with an oscillation detected by our method, a 3-way ANOVA with repeated measures yielded a significant main effect of eyes opening for alpha (F(1,50)=120.73, p<0.001) but not for SMR (F(1,18)=2.29, p=0.14) and a significant main effect for movement for SMR (F(1,19)=7.89, p=0.001) but not for alpha (F(1,25)=2.89, p=0.1). There is no main effect for the EEG device (percentage of time with alpha: F(1,25)=0.35, p=0.566, percentage of time with SMR: F=(1.33), p=0.26).

**Figure 7.**
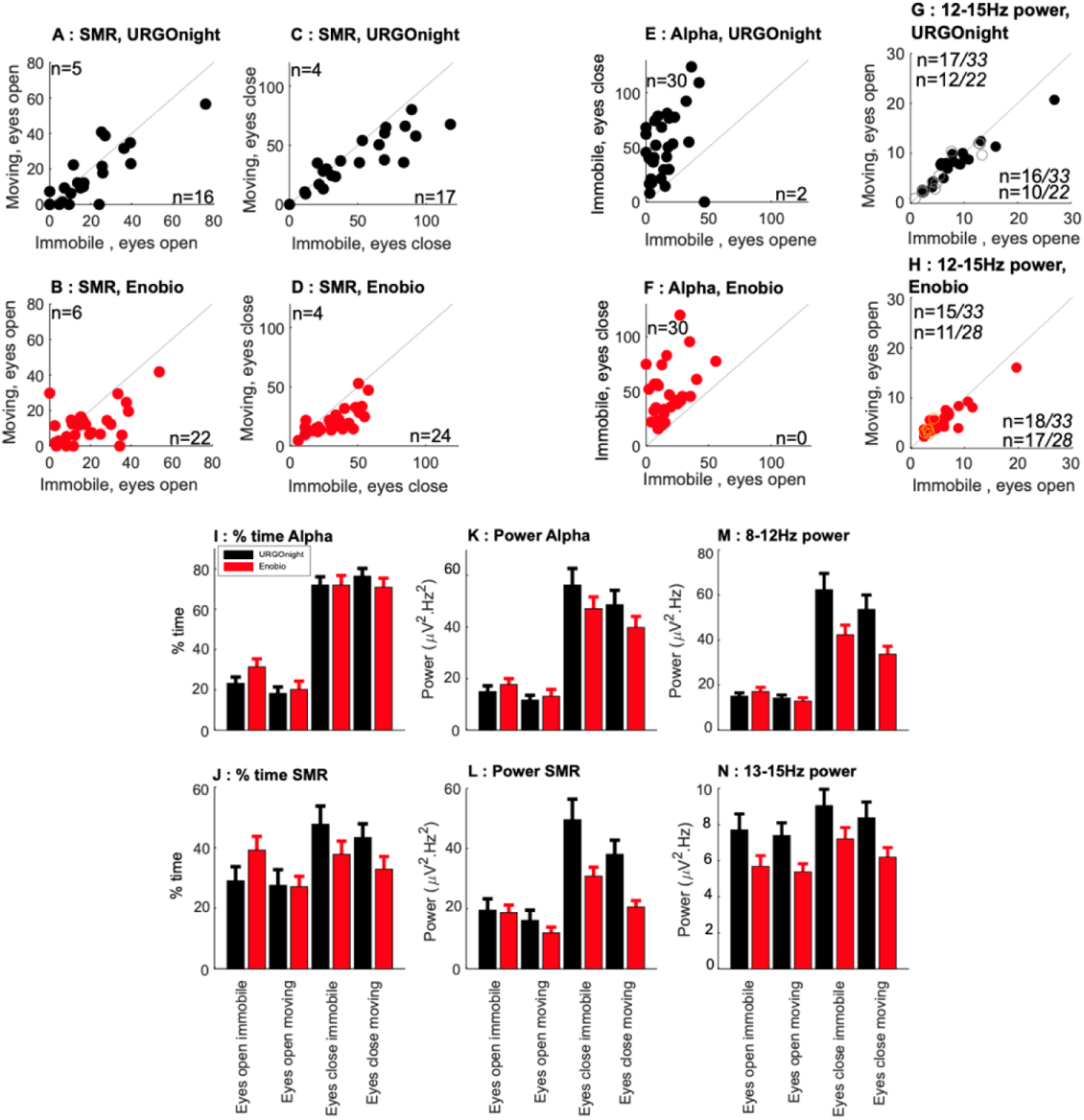
Evolution of the Alpha and SMR as a function of the dual blocking task. **A,B**. Correlation between the power of the detected SMR between mobile and immobile situations in subjects with open eyes. For all panels: URGOnight in *black* (A) and Enobio in *red (B)*. The number of subjects showing an increase or a decrease of the corresponding value is indicated on the figure. **C,D**. Same as A,B on subjects with eyes closed. **E,F**. Correlation between the power of the detected alpha between eyes open/closed situations in immobile subjects. **G,H**. Same as A;B but with the SMR defined as the average power in the 12Hz-15Hz frequency band. Grey and orange circles represent subjects for which the SMR was not detected with our method. **I,J**. Bar plot showing the percentage of time during which each rhythm is detected according to the dual blocking task (**I**: alpha, **J:** SMR). **K,L**. Average amplitude of the SMR and Alpha when the rhythm is detected according to the dual blocking task **(K:** alpha, **L:** SMR). M,N. Same as K,L but with the alpha defined as the average power in the 8-12Hz frequency band and the SMR defined as the average power in the 8-12Hz frequency band. The full statistics analysis of the panels 1-N is available in the supplementary materials.

The results obtained with the power of the two oscillations showed stronger effects. Indeed, power of both the alpha oscillations and the SMR is affected both by eye opening and by movement. This is likely due to the well-known effect of attention on both the alpha rhythm and the SMR, although one can also assume that the SMR is also contaminated by the alpha rhythm is the eyes closed condition. Nevertheless, we can see that the effect have a larger effect size for the congruent situations (Power alpha – dCohen : device 0.28, eyes 2.16, mov 0.70; % alpha – dCohen : device 0.1, eyes 1.86, mov 0.29; Power SMR - dCohen : device 0.74, eyes 1.31, mov 1.31; % SMR - dCohen : device 0.26, eyes 0.35, mov 0.64).

This is not the case when the two types of oscillation are defined according a to frequency range, especially for the SMR where the largest effect size is found foreye opening, and with only a small effect size for movement (12-15Hz - dCohen : device 0.68, eyes 0.71, mov 0.48; 8-12Hz - dCohen : device 0.58, eyes 1.48, mov 0.74).

Altogether, these results show that our detection method for alpha and the SMR is more efficient than the classical method that relies just on the average over a fixed frequency band. Finally, our results also show that URGONight is as efficient to detect t rue periodic components corresponding to alpha and the SMR as Enobio, a validated wireless medium-density EEG medical grade system.

## 4. Discussion

The URGOnight EEG headband has been developed to provide at-home SMR-based neurofeedback exercises to help users to improve sleep quality. Importantly, the efficiency of neurofeedback relies on the quality of the signal detected. To this day, many studies aimed at evaluating the signal quality in portable EEG devices by comparing them to conventional systems (Guger et al., 2012; Mathewson et al., 2017; Radüntz, 2018) or by comparing them to each other (Debener et al., 2012; Grummett et al., 2015; Kam et al., 2019; Zerafa et al., 2018). Nevertheless, most of these articles focused on practical application and did not investigate the quality of the device to detect a true oscillatory activity.

Poor headset placement could be one of the main limits for the at-home application in the neurofeedback systems used in laboratories. The choice of an adjustable headband with only two recording sites was made to improve the consistency in the positioning of the electrodes. The design of URGOnight is thus a trade-off between comfort and quality of the signal. Altogether, our results validate the quality of the EEG recordings obtained with URGOnight since it gives similar results as the one obtained with Enobio-20, one of the best portable EEG devices on the market. In addition, we provide a new method allowing the identification and the separation of the alpha and SMR with a single recording site C3/C4.

### 4.1. Quality of the EEG recordings obtained with URGOnight

The crucial step for a successful at-home neurofeedback system is to ensure the high quality of the signal recorded by a non-expert user. In this study, we have developed an automatic procedure of data selection that is fast enough to be done at the beginning of each recording session (less than a minute) and that ensures the quality of the signal (86% of the signals passes the criteria for the Enobio-20 and 80% for the URGOnight). Automatization of non-artefactual data selection could allow the users to train on good EEG signals at home.

We also showed that the EEG signal obtained with URGOnight is similar to Enobio-20, a wireless EEG device with wet electrodes that is used in clinical and fundamental research settings (Ratti et al., 2017). The response to SSVEP for stimulation frequency between 4Hz and 20Hz was similar for the two EEG devices. The main difference was the increase in power in the alpha band for URGOnight. We have shown that this increase is largely explained by the position of the reference electrode of URGOnight since we can reproduce this effect by a re-refencing procedure on an EEG database (64 channels) of 109 subjects recorded in eyes close situation (Goldberger et al., 2000; Schalk et al., 2004). We have demonstrated that an increase of power in the 8-15Hz in URGOnight range does not prevent the identification of both the alpha or SMR in most of the subjects. Nevertheless, increased alpha power recorded with URGOnight could explain some of the difference between alpha and SMR observed in this study. Kuhlman mentioned that bipolar derivation in the mediolateral plane showed little mu activity compared to the anteroposterior orientation (Kulhman, 1978). The position of the URGOnight reference close to PO7 is in the middle of these two orientations, and this might explain why SMR is slightly more difficult to detect with URGOnight compared to Enobio-20.

### 4.2. Problem of source identification for alpha and SMR with one site recording

During awake inactive states, excitation coming from the thalamus results in a desynchronized activity in the cortex which is captured by EEG recordings (Steriade and Llinas, 1988). This is associated with a blocking of occipital alpha rhythms during visual stimulation and with a blocking or desynchronization of central mu rhythms during somatosensory stimulation or movement (Berger, 1930; Chatrian et al., 1959; Gastaut, 1952). Some authors proposed the concept of sensory-specific alpha rhythms, in which the mu rhythm was viewed as a central “somatosensory alpha rhythm” in contrast to the occipital “visual alpha rhythm.” (Kulhman, 1978). Each ‘alpha’ rhythm would represent a deactivated or inhibited state of a specific primary cortex. Such inhibition (or passing into an ‘‘idling’’ state of the visual or motor area networks) can occur when attention is withdrawn from a specific modality, vision from the occipital alpha and motor movements for the mu rhythm (Niedermeyer, 1997; Pfurtscheller and Lopes da Silva, 1999). The frequency range of alpha rhythm is well defined between 8 and 12 Hz; however, the frequency band of the sensorimotor rhythm is less established. Initially the SMR was identified in cats within a frequency band of 12-15Hz. In humans, an analogue of this rhythm, often referred to as the mu rhythm, has been described but with frequency slightly lower ranging from 6Hz to 13Hz with an average of 9Hz (Kulhman, 1978). Despite such differences, when frequencies are considered, SMR is often used as synonym of mu rhythm even if SMR also includes a component in the beta band. This beta band is weaker and has different properties, which suggests a different neuronal origin (Cheyne, 2013).

In EEG studies, the separation between the SMR and alpha rhythm can be done by using their spatial localization: alpha is stronger over the occipital cortex while the SMR is maximal over the central/parietal cortex. General tools to perform the differentiation are based on multi-site recordings and include applying spatial filters, finding common spatial patterns or coherence between different recordings sites (Guger et al., 2000; McFarland et al., 1997; Schoppenhorst et al., 1980). In these studies authors do not deeply investigate oscillatory activity and do not mention whether true sensorimotor or visual-related oscillations are being captured by the method used. One of the reasons explaining the absence of careful comparison between the two rhythms is that the SMR is difficult to identify in most human subjects, despite a well-described mu-rhythm recorded from animals in the 12-15 Hz frequency band (Montaron et al., 1979; Philippens and Vanwersch, 2010) and in human with intracranial recordings (Arroyo et al., 1993). In fact, humans’ alpha oscillations prevent direct observation of the SMR due to significant overlap in their frequency bands. The situation is further complicated by the fact that the frequency bands of the SMR and alpha could significantly and independently vary across subjects (Kulhman, 1978).

Kulhman (1978) used the suppression of the SMR by movement and the suppression of alpha by eye opening to identify the two rhythms by contrast. Although mu activity was generally found in the same frequency range as the alpha rhythm, the two rhythms were separable in each subject in his study. By using EEG recordings from different sites on the scalp to differentiate the two rhythms, he found that rhythmic activity congruent with the functional characteristics of the occipital alpha rhythm was identified in 13 of the 14 subjects (93%), and rhythmic activity congruent with the functional characteristics of the mu rhythm was identified in 7 of the 14 subjects (50%).

In the present study, we employed the same behavioral procedure but adjusted for a device with one recording site in C3 (or C4). This method was initially developed by Donoghue and colleagues in order to separate periodic and non-periodic components and the EEG and thus to identify a true oscillation (Donoghue et al., 2020). We improved the speed of the computations so that it can be implemented in mobile and be utilized in real-time conditions for future neurofeedback application. Using such improved methodology, we identified the SMR and alpha rhythm in a reliable manner. Based on the evidence coming from the literature (Kulhman, 1978), the most frequently detected oscillations were defined as alpha, the second bump (if detected) corresponded to the SMR. We found an alpha-like rhythm in 32 out of 33 subjects with URGOnight and 30 out of 33 with Enobio, and we observed a blocking effect of eye opening in 30 subjects with URGOnight and 30 with Enobio, which corresponds to the identification of an unambiguous alpha rhythm in 90% of the subjects. We found an SMR-like oscillation in 22 subjects with URGOnight and 28 with Enobio, and we observed a blocking effect of active movement in 16 subjects with URGOnight and 22 with Enobio, which corresponds to the identification of an unambiguous SMR in 48%-66% of the subjects. This is much more than what has been found in early studies (Chatrian et al., 1959) but close to the proportions observed by Kuhlman (1978). It is worth pointing out that our detection method has the same efficiency with the data restricted to a one single recording site compared to a human expert analyzing multi-sites EEG obtained with recordings optimized for every subject to maximize the likelihood to detect the two rhythms (Kuhlman 1978).

### 4.3. Variation of alpha and SMR power with attention and commitment to the task

Our results show that the percentage of time during which we identify the SMR rhythm is more reliably associated with the blocking effect of movement than the SMR power that can be affected by eye closure as well. In other words, the movement condition will affect the period of time with the SMR but not with the alpha rhythm, although it may modify the amplitude of alpha. We also showed that the blocking effect of movement on SMR power was more visible in eyes closed compared to the eyes open condition. Two possibilities might explain this observation. First, the SMR might indeed be slightly more reliably detected in closed eyes situation. This is not supported by earlier studies that showed an antagonistic response of alpha and SMR: the focus on one modality decreased the attention towards the second modality (Kulhman, 1978). Moreover, direct intracranial recording of the mu rhythm in humans did not reveal any effect of eye closure on the oscillation but this study was qualitative and did not report any quantification of the power of the oscillation (Arroyo et al., 1993). Nevertheless, we can’t rule out the hypothesis that some subjects might improve their relaxation when having their eyes closed and that this condition would favor the emergence of the SMR. In that case, neurofeedback training would benefit (at least in some subjects) from being done with eyes closed.

The second possibility is that part of the SMR detected in eyes closed condition is contaminated by alpha rhythm and that the modifications observed are due to an indirect effect on alpha induced by the fluctuation of attention associated with the task. In fact, the power of both alpha and SMR is affected by attention (Anderson and Ding, 2011; Arroyo et al., 1993; Klimesch et al., 1998), although their link with attention and relaxation is complex (Glass and Kwiatkowski, 1970; Mundy-Castle, 1957). The interaction between the two tasks could therefore affect the power of both alpha and the SMR in a complex manner. During immobility, eye closure might be associated with movement imagination that will affect the power of SMR. Similarly, with eyes closed, the initiation of movement might be associated with a modification of attention that may lead to a decrease of the SMR rhythm. If the SMR is analyzed by averaging the 12-15Hz frequency band, it is not possible to dissociate the variation of the SMR-related oscillations from a leakage due to the modification of the alpha rhythm. Our method, which relies on the identification of a specific bump in the spectrogram, limits this bias although we cannot rule out the possibility that some of our effect s on the SMR are still due to leakage from alpha. This may explain why the amplitude of the SMR and alpha rhythm are modulated by both the eye opening and the movement conditions (although the effect size for the congruent situations are larger).

On the contrary, the periods during which the SMR and alpha are detected with our methods gives unambiguous results with SMR modulated by movement and alpha by eye opening. Altogether, we have shown that our method can be used to identify true oscillations that exhibit the properties of the SMR and that is separate from alpha oscillations.

### 4.4. Difference between alpha, SMR and 12-15Hz, specificity of neurofeedback training

In most neurofeedback studies, the SMR is defined as the average power between 12- 15Hz measured over central positions. There seems to be three reasons for this. First, the SMR was initially discovered in animal studies as visually distinct oscillations with a peak in the 12-15Hz frequency range (Brazier, 1963; Howe and Sterman, 1972). The mu rhythm in humans turned out to be an analogue of this rhythm but was detected at a lower frequency (Kulhman, 1978). Nevertheless, the 12-15Hz frequency band for the SMR was sometimes kept even in human studies. Second, the 12-15Hz was chosen for practical reason since it may contain a part of the SMR (by frequency leakage) while avoiding the frequency range of the occipital alpha that is the most prominent oscillation even in C3/C4 recordings. Finally, some neurofeedback studies using 12-15Hz as the SMR were successful both for enhancing this frequency band and for improving cognitive function or sleep (Cortoos et al., 2010; Gruzelier, 2014a, 2014b; Hoedlmoser et al., 2008; Schabus et al., 2014).

Nevertheless, success of 12-15Hz SMR-based neurofeedback is a still highly controversial topic (Fovet et al., 2017; Schabus, 2018, 2017; Schabus et al., 2017; Thibault et al., 2017; Witte et al., 2018). The demonstration of the specificity of SMR neurofeedback on a clinical use (choice of the right control group) and the optimization of the training parameters (coherent biomarker, feedback…) are the main controversial issues to overcome. In our study, we have clearly demonstrated that the power between 12 and 15Hz does not capture the properties of SMR (blocking effect induced by movement), on the contrary to our SMR detection method. Considering the possible overlap between SMR and alpha frequency bands, it is important to note that if the frequency of the SMR is higher than the frequency of the alpha rhythm then the 12-15Hz power might capture the alpha signal due to frequency leakage. On the contrary if the frequency of alpha is higher than the frequency of the SMR, then the 12-15Hz power will only capture some leakage of the alpha rhythm. This could be an explanation for the variable success rates of SMR modulation training found in the literature (Alkoby et al., 2018). In fact, the proportion of these two populations might vary in the neurofeedback studies (that contains a small number of subjects) and might well explain the success or failure of the 12-15Hz-based neurofeedback training. Importantly, our method will detect the SMR in every single subject no matter the frequency of the occipital alpha. It could be used in neurofeedback studies aiming at comparing feedback quality based on subject-specific SMR oscillation and on the 12-15Hz frequency band, which would be key to improve the efficacy of the procedure.

### 4.5. Importance of detection of an oscillation for SMR neurofeedback sleep therapy

In the seminal work of Sterman, SMR neurofeedback during wakefulness was found to improve sleep and increase the amount of sleep spindles (Sterman et al., 1970; Wyrwicka et al., 1962). In these studies, neurofeedback was done in cats for which SMR rhythm (a true visible oscillation in the signal) is in the exact same frequency range as sleep spindles. On the contrary, in humans, the SMR is observed at lower frequency (9-10Hz), while the sleep spindles are found as in cats between 10 and 15Hz. This raised the question of whether the positive effect found in some studies sleep using the power in the 12-15Hz frequency range is due to the enhancement of this specific frequency band or whether the positive effect is due to the enhancement of SMR in a subpopulation of subjects which SMR falls into the frequency range used for feedback.

To provide a more accurate measure of the SMR in terms of blocking effect induced by movement, our method defines the SMR as a true oscillation. Intracranial recording in humans showed that there is a true oscillation in the motor cortex which is difficult to observe in the EEG. This might be an interesting advancefor its use in neurofeedback-based therapy of sleep disorders. SMR neurofeedback is hypothesized to train the circuits associated with the generation of sleep spindles. Presumably, oscillations-based neurofeedback training is efficient only if the training focuses on already existing oscillations (Reichert et al., 2015). Thus, for a successful sleep improvement by the SMR-based neurofeedback, it is crucial to identify the specific SMR frequency band for each individual and to be able to distinguish it from the classical alpha oscillations.

To conclude, our study showed that URGOnight provides similar results compared to a validated portable device Enobio-20. Both alpha and SMR can be detected from the signal. Moreover, our SMR detection methods show a great improvement in capturing an oscillation affected by movement compared to the classical 12-15Hz frequency band. Thus, this method opens up research for a new type of SMR neurofeedback based on reinforcement of the SMR oscillation strictly speaking.

## Abbreviation

SMR: sensorimotor rhythm
IAF: individual Alpha frequency
SSVEP: Steady state visually evoked potential

## Ethical Approval

The experiments were carried out in accordance with the Declaration of Helsinki. The study was approved by the Committees of Protection of Persons (Comité de Protection des Personnes ”Est IV”), declared to the French National Agency for Medicines and Health Products Safety, and the study is registered on clinicaltrials.gov. Informed consent was signed by each participant before the experiment. Subjects received monetary compensation for their participation in the experiment.

## Consent to Participate

All participants provided informed consent.

## Code availability

All the codes of this study will be made freely available by email to the corresponding author. Any additional information required to reanalyze the data reported in this paper is available from the lead contact upon request.

## Authors’ contributions

RS performed the data acquisition, wrote matlab codes for data analysis and analyzed the data. KB coordinated the study, designed the study, performed data analyses and wrote the manuscript. ETC designed the study, provided valuable feedback during data analyzes and provided the URGOnight device. BS initiated this study. All authors contributed to, reviewed, provided feedback on, and approved the final manuscript.

## Acknowledgements

We would like to thank the time and effort put forward by our participants. We also thank Dmitri Bryzgalov, Thierry Gallopin, Sophie Bagur, Léa Lachaud, Charles Hernoux and Charles Verdonk for useful discussions. This study was funded by UrgoTech. The URGOnight device was provided by UrgoTech. We would also like to thank Robin Reynaud, Thomas Poujol and Magali Coquery for their help and tenacity across the development of the URGOnight device.

